# Liver angiocrine myeloid-derived growth factor protects against endothelial dysfunction in pulmonary arterial hypertension

**DOI:** 10.1101/2025.10.03.680334

**Authors:** Navneet Singh, Talia Volpicelli, Hongyang Pi, Sina Gharib, Elizabeth O. Harrington, Soban Umar, Peter J. Leary, Michael Fallon, Corey E. Ventetuolo, Olin D. Liang

## Abstract

Myeloid-derived growth factor (MYDGF) is a hepatic angiokine with protective effects in systemic vascular beds, but its role in pulmonary arterial hypertension (PAH) is unknown. We hypothesized that hepatic MYDGF deficiency contributes to pulmonary endothelial activation in PAH and that recombinant MYDGF could rescue endothelial injury. In the Sugen-hypoxia (SuHx) rat model, hepatic MYDGF expression was decreased, while pulmonary vascular cell adhesion molecule-1 (VCAM-1) expression was increased. Human hepatic sinusoidal endothelial cells exposed to pro-inflammatory macrophage conditioned media downregulated MYDGF, and recombinant MYDGF restored pulmonary artery endothelial cell resistance to inflammatory activation via MAP4K4–NFκB signaling. In the Brown University PHiNE PAH cohort (n=41 PAH, n=27 controls), plasma proteomics demonstrated increased MYDGF in PAH patients compared with controls, but MYDGF levels declined with worsening liver stiffness and correlated with higher pulmonary vascular resistance. In the independent Servetus PAH cohort (n=117), higher plasma MYDGF was associated with mortality and right ventricular dilation. Together, these findings demonstrate hepatic MYDGF deficiency in experimental PAH, tissue specificity of endothelial MYDGF to the liver, and MYDGF’s potential to mitigate pulmonary endothelial inflammation. However, human data suggest a paradoxical association of elevated circulating MYDGF with adverse outcomes, underscoring the complex biology of angiogenic growth factors in PAH. MYDGF may represent a novel hepatic angiokine linking systemic inflammation, liver dysfunction, and pulmonary vascular disease.

## Introduction

Circulating angiogenic growth factors have an established but complex role in the pathobiology of pulmonary arterial hypertension (PAH). Myeloid-derived growth factor (MYDGF) is a hepatic growth factor with angiogenic properties produced primarily by cells of myeloid origin, particularly macrophages, as well as hepatocytes and hepatic endothelial cells (ECs)(1). MYDGF confers protection in systemic vascular beds including the liver, heart, and kidney (2), but its role in pulmonary vascular disease is unknown. We examined whether MYDGF is implicated in PAH. We hypothesized that hepatic MYDGF is deficient or ineffective in PAH and that *in vitro* MYDGF supplementation can protect against inflammatory EC activation. In two independent PAH cohorts without known liver disease, we asked whether circulating MYDF is associated with clinical measures.

## Methods

The Sugen-hypoxia (SuHx) rat model was established as previously described (5). The left lobe of the liver and lung from each animal was fixed, sectioned and stained for MYDGF (Thermo MA5-31965). Liver tissue was homogenized in RIPA (Thermo) and TRIzol (Thermo) for protein and RNA, respectively. Study staff were blinded. Protein underwent immunoblot for MYDGF and qPCR per protocol. Commercial human hepatic sinusoidal ECs (HSECs) and pulmonary artery ECs (PAECs) (Lonza) were cultured with and without M1 macrophage pro-inflammatory conditioned media (M1CM) which was prepared as previously described (6) and with or without 100nM recombinant MYDGF (R&D 10231-MY).

Plasma was obtained from a prospective observational cohort of PAH and control participants recruited from the Center for Advanced Lung Care at Brown University Health. Forty-one PAH and 27 controls were included. The medical record was reviewed to collect clinical data and non-invasive measures of liver dysfunction were calculated including the fibrosis 4 (FIB-4) score; patients were stratified into low (< 1.3) and intermediate/high (≥1.3) scores based on accepted thresholds (3). Plasma was submitted for data-independent acquisition (DIA) LC-MS/MS. Relative protein abundances were calculated by normalizing each protein’s intensity to the total protein signal per sample. The Seattle Right Ventricle Translational Science (Servetus) cohort is a prospective observational PAH cohort recruited from the University of Washington (4). Plasma proteomics were performed via the aptamer-based SomaLogic, Inc platform (Boulder, CO). The study was approved by the Institutional Review Boards (IRBs) at Brown University Health (IRB# 2221835) and the University of Washington (IRB# STUDY00003387).

Statistical analysis was performed in GraphPad Prism (v10.4.1) and RStudio (v4.4.1). Student’s t-test was used to compare experimental data. Pearson’s correlation or logistic regression was used to compare MYDGF expression with clinical outcomes. A p<0.05 was considered statistically significant.

## Results

SuHx rat livers had decreased MYDGF staining as compared to controls (6.0 % vs 10.4%, p=0.001), though expression was retained along hepatic sinusoid walls, the site of sinusoidal ECs. In the SuHx rats with decreased hepatic MYDGF, VCAM-1 expression was increased along the pulmonary endothelium (61.3% vs 31.4%, p=0.001) (**Figure 1A**). Immunoblot analysis and qPCR of homogenized liver tissue confirmed less MYDGF protein (mean 1.6, range 1.0-2.1 vs 1.9, range 1.0-2.8; p=0.23) and mRNA (mean 0.14, range 0.13-0.15 vs 0.18, range 0.12-0.23; p=0.14) in SuHx rat livers compared to controls but did not reach statistical significance. Only scant MYDGF expression could be detected in SuHx lung ECs (**Figure 1B**). HSECs cultured in M1CM for 72 hours had decreased MYDGF protein expression as compared to controls (0.19 vs 0.77, p=0.07)(**Figure 1C**). Phosphorylated MAP4K4 and NFκB (p65) expression was increased in PAECs cultured in M1CM as compared to controls; this could be rescued by treatment with 100nM recombinant MYDGF (**Figure 1D**).

**Figure 1.**
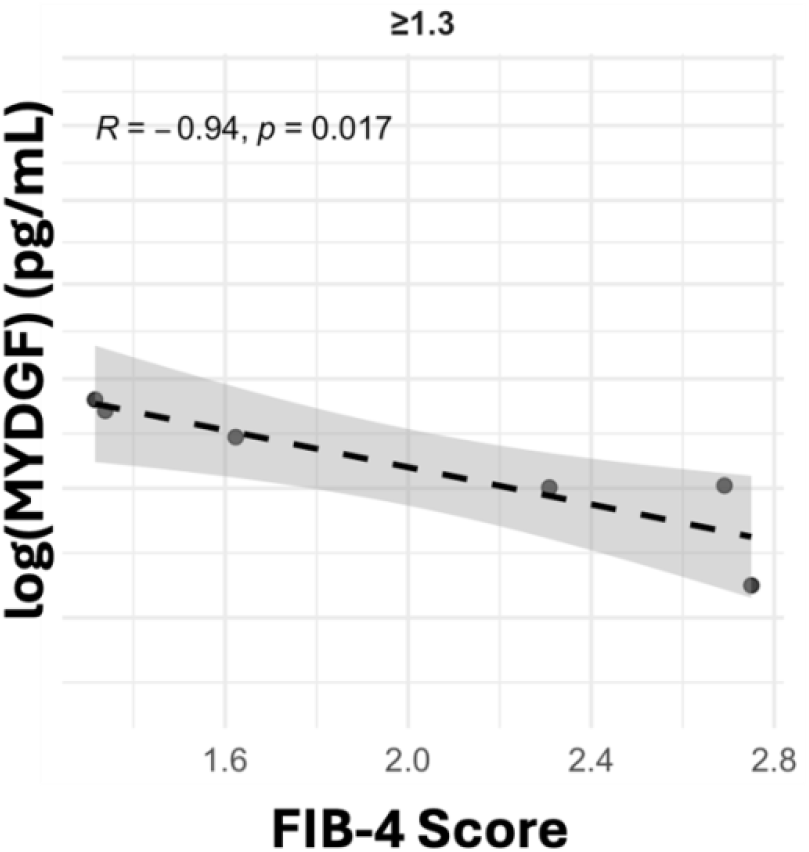
Plasma MYDGF decreases as liver stiffness (FIB-4 score) increases. Pearson correlation was performed to evaluate the association of plasma MYDGF from PHiNE subjects and liver stiffness as measured by the validated non-invasive *FIB-4 score. MYDGF=myeloid derived growth factor. FIB-4=fibrosis 4. PHiNE=Pulmonary Hypertension in New England*.

In the Brown cohort, a total of 1,271 proteins were differentially expressed between PAH and control subjects. MYDGF was detected in 21 subjects (n=13 PAH and n=8 controls). Normalized MDYGF expression was increased by 13% in PAH vs control (log_2_(ratio)=0.20, p=0.0007). In subjects with intermediate/high FIB-4 scores (i.e., increased liver stiffness), MYDGF levels decreased as liver stiffness increased (R=-0.94, p=0.02)(**Figure 2**). Pulmonary vascular resistance (PVR) was higher in these subjects compared to those with low FIB-4 scores (median (IQR) 6.7 (3.1-9.8) vs 3.6 (2.5-5.9) Wood units, p=0.07). The Servetus cohort included 117 PAH patients (4). Logistic regression of dichotomized plasma MYDGF levels adjusted for age, sex, BMI and PAH etiology demonstrated that high MYDGF levels (above the median) were associated with a lower likelihood of survival (ß -1.58, p=0.009) and right ventricular dilation (ß=-0.31, p=0.002)

**Figure 2.**
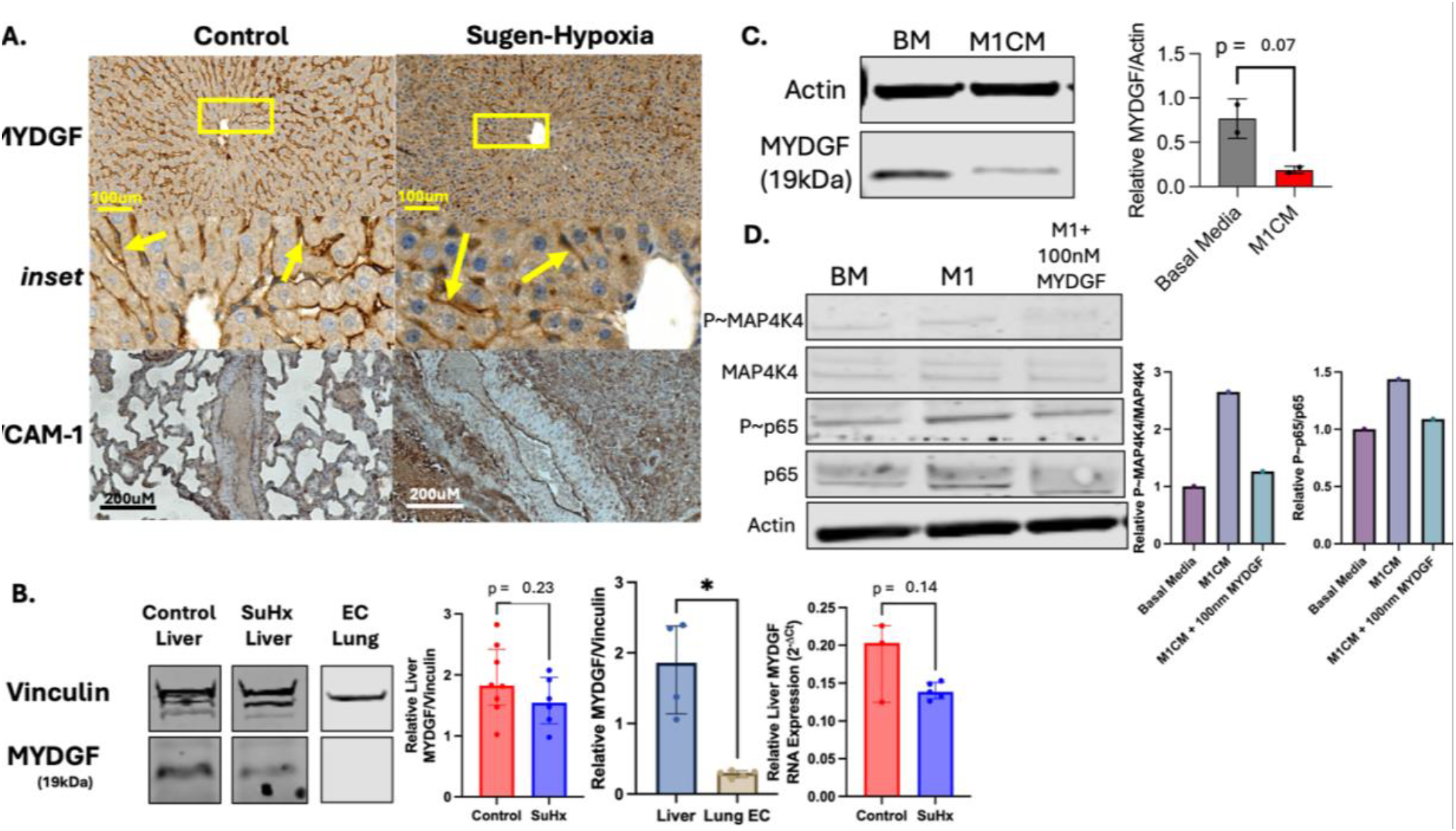
Hepatic MYDGF Deficiency Links Inflammation to Pulmonary Endothelial Injury. **A**. Representative liver sections of control (left) and SuHx (right) rats stained for MYDGF, and lung sections stained for VCAM-1 (brown). Despite decreased overall MYDGF staining in SuHx liver, expression is retained in HSECs (arrows).**B**. Quantification shows decreased MYDGF protein and mRNA expression in SuHx livers vs controls; no MYDGF could be detected in SuHx lung endothelial cells suggesting tissue specificity of MYDGF in this model. **C**. Commercial human HSECs exposed to M1 pro-inflammatory media (M1CM) have decreased MYDGF protein expression vs basal media. **D**. Commercial human PAECs have decreased pro-inflammatory activation (phospho-MAP4K4 and phospho-p65) when rescued with 100nM MYDGF. Graphs represent phospho-protein normalized to actin and relative to total MAP4K4 or p65, respectively. *SuHx=Sugen-Hypoxia. MYDGF=Myeloid-Derived Growth Factor. HSEC=hepatic sinusoidal endothelial cell. PAEC=pulmonary artery endothelial cell. BM=basal media. M1CM=M1 pro-inflammatory conditioned media*. All images taken with 10x objective; scale bar represents 200μM. n=2-3 rats in each group.

## Discussion

This is the first demonstration that PAH is characterized by hepatic MYDGF deficiency and that MYDGF can protect the pulmonary endothelium from inflammatory activation. In two independent PAH cohorts without known liver disease, plasma MYDGF levels were linked to PAH outcomes and subclinical liver stiffness. Although we demonstrated evidence of MYDGF’s protective potential *in vivo* and *in vitro*, our human data suggests that higher MYDGF levels are linked to mortality. This discrepancy recapitulates prior work on hepatic growth factor (12) and speaks to the complex biology of angiogenic growth factors We propose that myeloid-derived MYDGF is increased as part of the inflammatory activation of PAH, which in turn disrupts hepatic secretion of MYDGF into the circulation, thereby explaining some of the directionally inconsistent results from PAH patients. In fact, hepatic MYDGF was decreased and localized along hepatic sinusoidal walls, the location of sinusoidal ECs, a site of liver homeostasis and regulation of blood flow, filtration and immune functions (7). There was virtually no endogenous MYDGF expression noted in the pulmonary endothelium, suggesting tissue specificity of endothelial MYDGF expression for the liver. HSECs cultured in pro-inflammatory media recapitulated this decreased expression and treatment with recombinant MYDGF rescued PAECs from pro-inflammatory activation. Together, these results suggest MYDGF is a novel hepatic angiokine that modulates pulmonary vascular disease.

MYDGF is an angiogenic growth factor which appears to play a protective role in systemic inflammatory and fibrotic injury. There is evidence that MYDGF promotes EC repair in acute lung injury (8) and counterbalances pro-fibrotic influences from TGFβ1 (9), a member of the superfamily to which BMP signaling belongs and a central pathway in PAH. In rodent models, MYDGF acts in an endocrine fashion to reduce coronary inflammation, atherosclerosis and endothelial damage via the MAP4K4-NFκB pathway (10). MYDGF is primarily produced by myeloid-lineage cells however the importance of the protein’s production by the liver was recently discovered (1). Here, we confirm the biological importance of hepatic MYDGF by demonstrating its deficiency in the livers of a leading animal model of PAH and that increasing liver stiffness in humans is linked to decreased plasma protein. Our observation that a pro-inflammatory milieu can promote decreased MYDGF expression in HSECs provides a potential mechanistic connection to the systemic inflammation that occurs in PAH (11) and reinforces our hypothesis that the liver can potentiate pulmonary vasculopathy in PAH via interorgan crosstalk.

Although novel, our observations require confirmation using *in vivo* experimental models and prospective human studies with larger sample sizes. Additional work is needed to parse the contributions of myeloid- vs hepatic-MYDGF. Specific mechanisms of MYDGF’s protection against pulmonary EC inflammation need to be explored and confirmed.

In summary, we offer the first observations of hepatic MYDGF deficiency in both experimental and human PAH, its tissue specificity in the liver and evidence supporting its potential as a protective therapeutic against pulmonary endothelial injury.

